# Reflection Knockoffs via Householder Reflection: Applications in Proteomics and Genetic Fine Mapping

**DOI:** 10.1101/2025.01.16.633369

**Authors:** Yongtao Guan, Daniel Levy

**Affiliations:** National Heart, Lung, and Blood Institute

**Keywords:** Knockoff filter, reflection knockoff (ReKo), proteomics, fine mapping

## Abstract

We present a novel knockoff construction method, and demonstrate its superior performance in two applications: identifying proteomic signatures of age and genetic fine mapping. Both applications involve datasets of highly correlated features, but they differ in the abundance of driver associations. Our primary contribution is the invention of the reflection knockoff, which is constructed from mirror images – obtained via Householder reflection – of the original features. The reflection knockoffs substantially outperform Model-X knockoffs in feature selection, particularly when features are highly correlated. Our secondary contribution is a simple method to aggregate multiple sets of identically distributed knockoff statistics to improve the consistency of knockoff filters. In the study of proteomic signatures of age, single feature tests showed overly abundant proteomic association with age. Knockoff filters using reflection knockoffs and aggregation, however, revealed that a majority of these associations are hitchhikers instead of drivers. When applied to genetic fine mapping, knockoff filters using reflection knockoffs and aggregation outperform a state-of-the-art method. We discuss a potentially exciting application of reflection knockoffs: sharing genetic data without raising concerns about privacy and regulatory violations.

## 1 Introduction

Scientific research often deals with datasets containing strongly correlated features. One example is proteomics data, in which high correlation can be attributed to several factors including the inherent correlations among multiple proteins, protein-protein interactions, and the imperfect specificity of the assay. Another example is genetic fine mapping, where single nucleotide polymorphisms (SNPs) in a locus may be highly correlated due to linkage disequilibrium. For these types of highly correlated data, identifying features that are truly associated with a phenotype of interest is a critical but challenging task. One key difficulty is controlling for false discovery, i.e., “hitchhikers” that show association through correlation with genuinely associated “driver” features. The knockoff filter (Barber and Candés, 2015; Candés et al., 2018) is a statistical framework designed to address this challenge by providing a robust way to identify true associations while controlling the false discovery rate (FDR).

Knockoff filters consist of four components: 1) constructing knockoff features; 2) conducting association analysis by jointly analyzing the original features and their knockoffs. The joint analysis can be conducted with either glmnet (Friedman et al., 2010), which implements penalized regression methods, including LASSO (Tibshirani, 1996) and Elastic-Net (Zou and Hastie, 2005), or fastBVSR (Zhou and Guan, 2019), which implements an iterative complex factorization algorithm to fit Bayesian variable selection regression (Guan and Stephens, 2011), or with other machine learning algorithms, including support vector machines (Cortes and Vapnik, 1995) and random forests (Breiman, 2001); 3) computing *knockoff statistics W* : a vector of differences in the feature importance measure between the original features and their knockoffs, where the feature importance measure includes the magnitude of regression coefficients (Beta) and posterior inclusion probability (PIP); and 4) determining the knockoff threshold from *W* at a nominal FDR level (e.g.,10%) to select features.

The most important component of the knockoff filter is constructing knockoffs 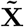 conditioning on the original features **X**. This is also the focus of our work. For the knockoffs to have the desired property of controlling FDR, **X** and 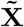 must follow:

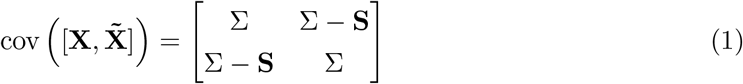

where **S** *≥* **0** is a diagonal matrix that satisfies

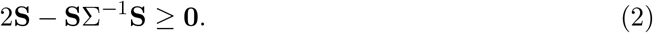

**S** controls the degree of uncorrelated-ness between **X**_*j*_ and 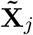, and larger **S** values bring more power to the knockoff filter. Another key condition for 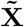 is that it must satisfy that, conditional on 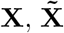 is independent of phenotypes of interest, which is automatically satisfied if the construction of 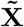 is agnostic to the phenotypes.

A critical component in generating both Fixed-X and Model-X knockoffs is estimating **S**. Three methods have been developed to estimate **S**: 1) The equi-correlated method where **S** is a scaling of an identity matrix; 2) The SDP method where **S** is obtained by optimizing a natural objective function subject to the inherent constraint using a semidefinite program (SDP) (Barber and Candés, 2015); and 3) A two-stage method where the first stage produces an SDP-estimate of **S**, but using an approximated feature covariance matrix, and the second stage linearly scales the SDP-estimated **S** (Candés et al., 2018). The SDP estimated **S** is more powerful than **S** estimated using the other two methods.

In this paper, we introduce a novel method to estimate **S** by first constructing mirror images **Y** of the original feature matrix **X** via Householder reflection. The difference, 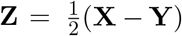, is orthogonal to **X** and thus **T** = cov(**X, Z**) is a diagonal matrix, where cov(**X, Z**) denotes a sample covariance matrix with **T**_*jk*_ = cov(**X**_*j*_, **Z**_*k*_) denoting sample covariance between two column vectors **X**_*j*_ and **Z**_*k*_. We define **S** = *α***T**, and estimate the scalar 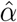 by examining the largest eigenvalues of the scaled **Z**. Such 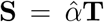 leads to more powerful knockoffs. We call the knockoffs constructed via Householder reflection *reflection knockoffs* (ReKo). Since its optimality comes from the leading eigenvalue and its computation does not require optimization, ReKo is numerically stable.

Due to the inherent randomness in constructing knockoffs, the knockoff filter may produce inconsistent results when applied to different copies of knockoffs. A second contribution of the paper is a simple method we propose to aggregate multiple sets of identically distributed knockoff statistics: pooling multiple knockoff statistics to determine the knockoff threshold and selecting features by comparing the average knockoff statistics with the knockoff threshold. This aggregation method produces more consistent results in real data analysis.

We apply reflection knockoffs and aggregate multiple sets of knockoff statistics to study two outstanding problems in genetics and proteomics, where the complex patterns of high correlation among features pose a challenge for existing methods. In the study of the proteomics signature of age, proteomic features appear to be overly abundant in association with age. Our study using reflection knockoffs revealed that the majority of single-feature associations observed between proteomic features and age are hitchhikers. In genetic fine mapping linkage, disequilibrium between nearby genetic variants makes it difficult to distinguish driver signals from hitchhikers. We demonstrate that reflection knockoffs significantly improve the power of knockoff filters for identifying driver variants.

## 2 Results

### 2.1 Reflection Knockoffs

A key observation leading to our invention of reflection knockoffs is that the reflection of a feature **X**_*j*_ over the hyperplane spanned by other features **X**_,−*j*_ produces a **Y**_*j*_ that is an almost perfect knockoff for that feature. Such **Y**_*j*_ can be computed using Householder reflection. Indeed, cov(**X**_*j*_, **Y**_*k*_) = cov(**X**_*j*_, **X**_*k*_) for *j* ≠*k* (Lemma 1 in Materials and Methods), where cov denotes sample covariance. As a candidate for knockoff of **X**, a linear combination of **X** and **Y** has the correct sample covariances between different column vectors, but not correct sample variances. The variance, however, can be fixed by adding a term that is uncorrelated with columns of **X** and **Y**, and this is the essence of a formulation provided by Barber and Candés (2015).

The Householder reflection regresses **X**_*j*_ on **X**_,−*j*_ to obtain a fitted value 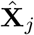, then computes the reflection 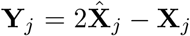. The residual of regression is 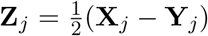. Therefore **Z**_*j*_ is orthogonal to 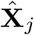 and uncorrelated with all other features. Thus, the sample covariance matrix **T** = cov(**X, Z**), where **T**_*jk*_ = cov(**X**_*j*_, **Z**_*k*_), is a diagonal matrix with **T** *≥* **0** (Materials and Methods). We define **S** = *α***T** and seek the largest *α* such that **S** satisfies the constraint in Equation (2). The optimal 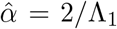, where Λ_1_ is the leading eigenvalue of the sample covariance matrix **C** = cov(**ZT**^−1*/*2^), where **C**_*jk*_ is the sample covariance between *j*-th and *k*-th column of matrix **ZT**^−1*/*2^. The knockoffs can be computed following Equation (8) in Materials and Methods. Notably, constructing reflection knockoffs requires no optimization and is numerically stable.

For the knockoff to have power, we need 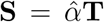 to be bounded away from 0. Let’s examine 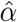 and **T** in turn. Numerically cov(**ZT**^−1*/*2^) is similar to an identity matrix, with diagonal entries equal to 1 and small off-diagonal entries. By Lemma 1 we have var(**Z**_*j*_) = **T**_*jj*_ where var denote sample variance. Since **T** > **0** is diagonal, we have diag(cov(**ZT**^−1*/*2^)) = diag(**I**_*p*_). The off-diagonal element cov(**ZT**^−1*/*2^)_*jk*_ is the sample correlation between residuals **Z**_*j*_ and **Z**_*k*_. Assuming cov(**X**_*j*_, **X**_*k*_) = *r*, and that both are uncorrelated with other features, a straightforward computation yields cov(**Z**_*j*_, **Z**_*k*_) = *r*(*r*^2^ − 1). Therefore both large (close to 1 or −1) and small (close to 0) correlations produce small cov(**Z**_*j*_, **Z**_*k*_). This calculation also applies when *r* is the conditional correlation, which can be obtained from the sample precision matrix cov(**X**)^−1^. When features are correlated, the conditional correlation between two features tends to be small. Then by the Gershgorin circle theorem, the eigenvalues satisfy |Λ_*j*_ − 1| *≤* ∑_*k* ≠ *j*_ |cov(**ZT**^−1*/*2^)_*jk*_|. So as long as the off diagonal entries are small and linearly diminishing with the dimension of cov(**ZT**^−1*/*2^), the leading eigenvalue Λ_1_ will be small, and 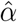 will be bounded away from 0.

By **T**_*jj*_ = var(**Z**_*j*_), for the reflection knockoff to have power the size of residual **Z**_*j*_ needs to be bounded away from 0. When features are highly correlated, however, var(**Z**_*j*_) tends to be small, as **X**_*j*_ is well explained by **X**_,−*j*_. Introducing shrinkage priors via ghost samples (Materials and Methods) proves to be an effective way to keep var(**Z**_*j*_) away from 0. This is a rare example where both sides of the bias and variance tradeoff are beneficial – we have more consistent estimates of **Z** while keeping the size of **Z** away from 0. Intuitively, when the mirror image is too close to the original feature, the priors bend the mirror so that the image appears farther away in the mirror without introducing obvious distortion. The ghost samples also enable reflection knockoffs to be constructed when the number of features is larger than the number of samples, significantly broadening the scope of applications.

### 2.2 Combined knockoff filter

We designated a knockoff filter by how the knockoffs were constructed, how the associations were analyzed (either using statistical methods or software packages that implement them), and what feature importance measure was used. For example, Model-X⊗glmnet⊗Beta means a knockoff filter that constructs Model-X knockoffs, uses glmnet (a R package) to conduct joint association analysis, and chooses the regression coefficient as the importance measure; ReKo⊗fastBVSR⊗PIP means a knockoff filter that constructs reflection knockoffs, performs association analysis using fastBVSR, and uses PIP (posterior inclusion probability) as the importance measure.

When comparing two knockoffs with everything else being equal, we designate knockoff filters in a manner such as⊗glment⊗Beta. Thus, by saying we are comparing Model-X and ReKo based on ⊗glment⊗Beta, we mean comparing power and type I errors between the knockoff filters Model-X⊗glmnet⊗Beta000000000000000000000000000000000 and ReKo⊗glment⊗Beta. We may omit the importance measure from the notation of knockoff filters when it is clear from the context. For example, ⊗glment means ⊗glment⊗Beta, and ⊗fastBVSR means ⊗fastBVSR⊗PIP.

A *combined knockoff filter ⊗*combined combines results from knockoff filters ⊗glmnet and ⊗fastBVSR, such that a feature is selected as positive by the combined filter if and only if it is selected by both ⊗glmnet and ⊗fastBVSR. The combined knockoff filter provides a means to combine results from Frequentist and Bayesian inferences.

### 2.3 Aggregating multiple knockoff statistics

Due to random component in their construction, knockoffs are inherently random. Consequently, inferences based on a single knockoff statistics can be inconsistent. Generating multiple knockoffs and combining multiple knockoff statistics can effectively increase consistency and power of a knockoff filter (Gimenez and Zou, 2019; He et al., 2021a). The existing methods that combine multiple knockoffs assume that feature importance measures can be obtained for each copy of knockoffs independently of the original features (He et al., 2021a), while for our target applications, it is important to jointly fit a model that contains both origin features and a copy of their knockoffs. We developed a new method that combines multiple sets of knockoff statistics (Material and Method).

Our aggregating method concatenates multiple sets of knockoff statistics to identify a knockoff threshold at a nominal FDR level, and calls positives by comparing average knockoff statistics of each feature with the knockoff threshold. Intuitively, concatenation increases the sample size used to estimate thresholds and provides a more stable empirical estimate of the distribution of null and non-null features. Concatenation reduces variance in the estimation of threshold-relevant ratios, making the threshold estimation more stable while preserving the original FDR control. Under the null hypothesis, if *m*-th knockoff statistic of *j*-th feature 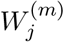 is symmetric around zero for each *m*, their average 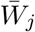 remains symmetric around zero. Consequently, 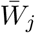 retains the crucial null property necessary for FDR control. Averaging also reduces the variance of knockoff statistics for each feature, providing more stable and reproducible feature selection while maintaining theoretical FDR guarantees. Figure 1 provides an example to demonstrate that aggregating multiple copies of knockoffs increases power. For the numerical analysis conducted in this paper, all results were obtained by aggregating 10 knockoff statistics, unless otherwise noted.

**Figure 1.**
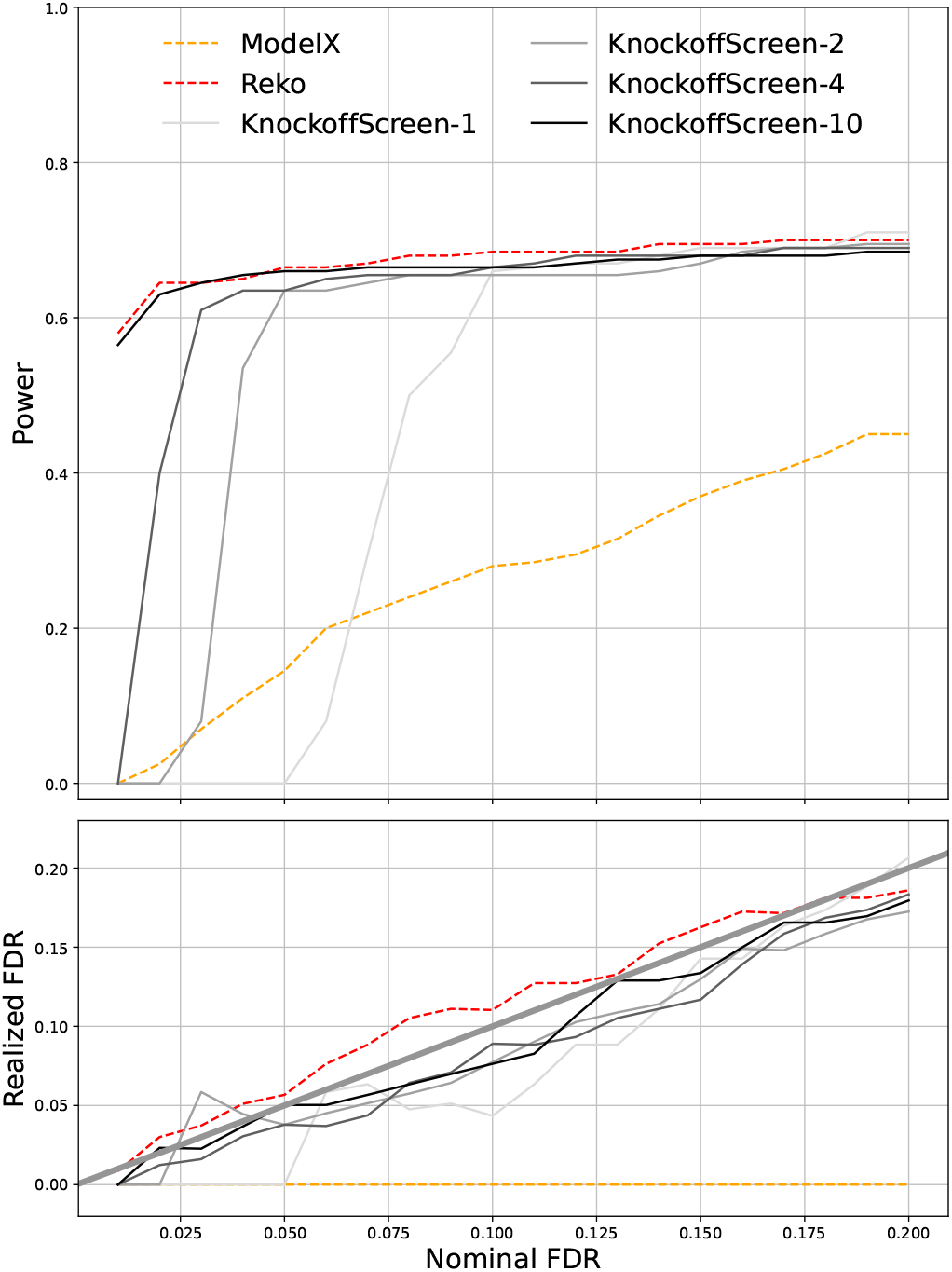
Aggregating knockoff statistics improve power. Top panel is nominal FDR vs power; Bottom panel is nominal FDR vs realized FDR. The knockoff filter used to call positives is ⊗glmnet for all. Results of Model-X and ReKo were obtained by aggregating 10 copies of knockoff statistics. For KnockoffScreen, we produced results by using a single copy of knockoff statistics and by aggregating 2, 4, and 10 copies of knockoff statistics. The results are based on simulated features with *n* = 2000 and *p* = 800. More details of simulation can be found in Supplementary Section 2.

### 2.4 Comparing ReKo with other knockoffs

We first compared ReKo with Fixed-X and Model-X knockoffs using synthetic features (Supplementary Tab. S2). Building upon knowledge gained, we then focus on comparing ReKo with Model-X and KnockoffScreen (He et al., 2021a) using synthetic features of various simulation parameters. These include high and low content of association signals (Supplementary Fig. S2 and S3.), different levels of feature correlations (Supplementary Fig. S4), and non-normal feature distributions (Supplementary Fig. S5). With synthetic features, we observed that ReKo is on par or slightly better than KnockoffScreen, and both are noticeably outperform Model-X. Since synthetic feature may not capture intricate correlation in the real datasets, we use real features for comparison.

The first real dataset is a subset of 1459 proteomic features from 9914 UK Biobank samples. The largest correlation between different features is 0.98. We simulated phenotypes using 200 randomly selected features as drivers. Their effect sizes were independently drawn from the standard normal distribution, but rescaled so that the PVE, or Proportion of Variance Explained, was 0.90. The large PVE used here was motivated by the strong signals observed in real phenotypes. Power and realized FDR were computed by pooling results from 100 replicates. For each replicate, we aggregated knockoff statistics from 10 copies of knockoffs (independently generated for each replicate) to call true and false positives at nominal FDR levels ranging from 0.01 to 0.20.

Figure 2 presents the results, and we make the following observations: 1) Model-X appears to be overly conservative on the realized FDR, and its power suffers consequently. 2) Both ReKo and KnockoffScreen showed inflated realized FDR for knockoff filters ⊗glmnet and⊗fastBVSR. With ⊗combined, however, both ReKo and KnockoffScreen have their realized FDR under control. 3) ReKo performs on par with KnockoffScreen, the slightly better power observed is likely due to ReKo’s slightly higher realized FDR. 4) Knockoff filter ⊗fastBVSR appear to be more powerful than ⊗glmnet. This is particularly true for Model-X, whose realized FDRs are under control for both knockoff filters. This echoes our prior studies (Guan and Stephens, 2011; Qi et al., 2018) that demonstrated advantage of Bayesian variable selection over panelized regression.

**Figure 2.**
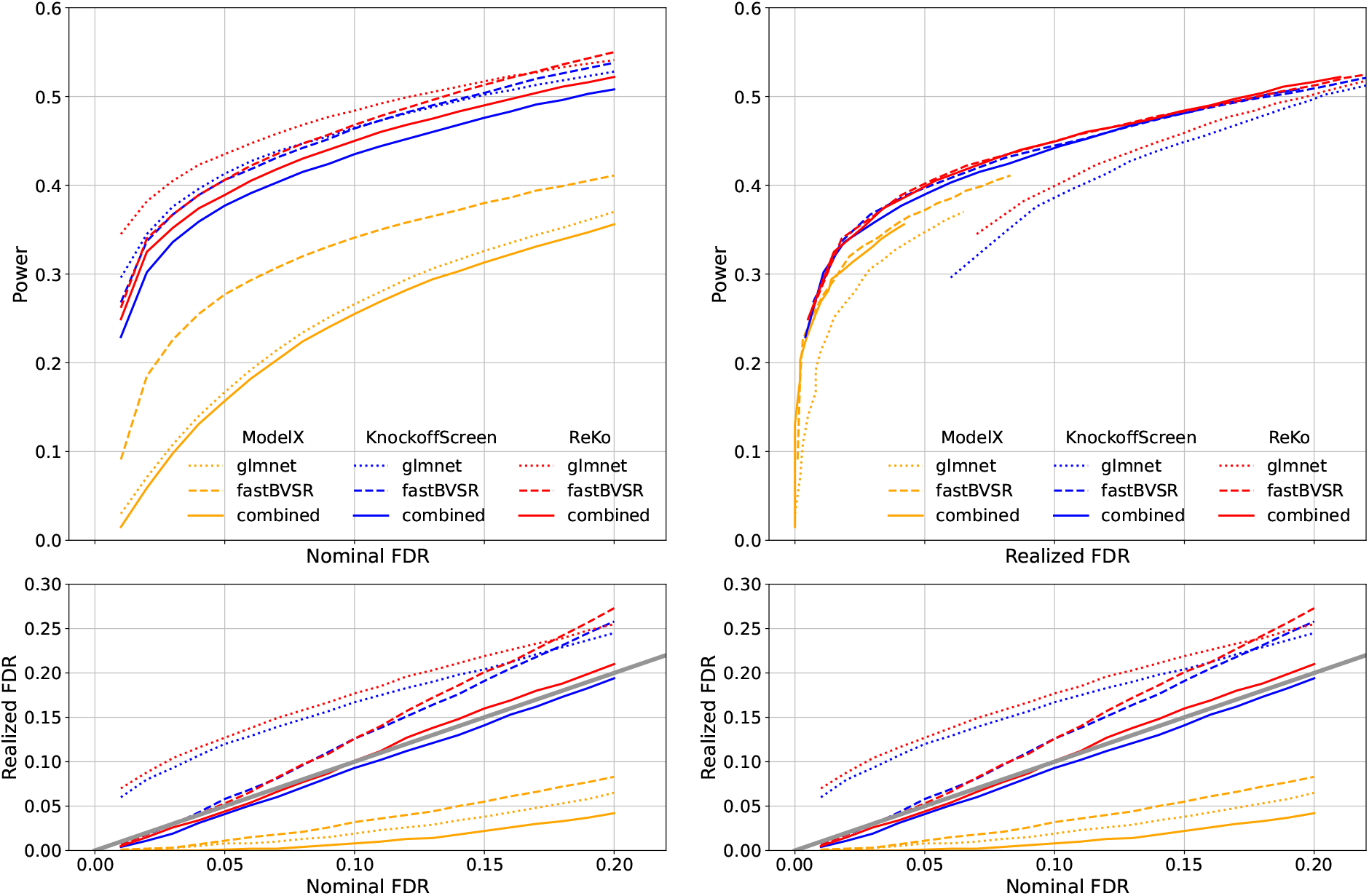
Power and realized FDR comparison between different knockoffs. The features are proteomics data from UK Biobank.. Left panel: the top plot is *nominal* FDR vs. power. Right panel: the top plot is *realized* FDR vs. power. The bottom plots of two panels are identical. Different line styles represent different knockoff filters and different colors represent different knockoff methods. The power and realized FDR were computed from 100 replicates, and each replicate aggregates 10 knockoff statistics.

The second dataset of real features consists of genotypes. And our goal is to investigate how high correlation affects performance of knockoff filters. From a trio-phased dataset (Guan and Levy, 2024a) we extracted features of paternal and maternal alleles from genomic region *HLA-DQB1*, which is known for its strong and complex pattern of linkage disequilibrium. We extracted three sets of features each having a different threshold for highest correlation among features: 0.90, 0.95, and 0.99. This is achieved by progressively thinning features in such a way that if a pair of features has a correlation that is higher than a threshold, we randomly remove one feature. To simulate phenotypes we randomly select 8 causal features, simulate each effect size with standard normal but rescale the effect sizes so that the PVE is 0.20. For each simulated phenotype, we independently generated knockoffs using Model-X, KnockoffScreen and ReKo.

Since this simulation is designed to illustrate the application of knockoff filters to study fine mapping, we compared results with SuSiE (Wang et al., 2020), a Bayesian method for fine mapping. We used its PIP outputs to call positives for SuSiE: at a nominal FDR *α*, a feature is called positive if its PIP is greater than 1 − *α*.

Figure 3 left panel summarizes the results for KnockoffScreen⊗combined, ReKo⊗combined, and SuSiE. We make the following observations: 1) At correlation thresholds 0.90 and 0.95, both ReKo⊗combined and KnockoffScreen⊗combined appear to have well-controlled type I error. 2) ReKo appears to have more power than KnockoffScreen, and at less stringent nominal FDR, ReKo appears to have more power than SuSiE. 3) At correlation 0.99 both ReKo⊗combined and Knockoff⊗combined show inflated realized FDR. 4) As the correlation threshold increases, the power of all methods decreases. 5) SuSiE has almost flat realized FDR vs. nominal FDR, suggesting that the calibration of its PIP remains challenging in real data analysis, an observation also made by others (**?**). But its power increases with nominal FDR, and at a more stringent nominal FDR, SuSiE is more powerful than both ReKo and KnockoffScreen.

**Figure 3.**
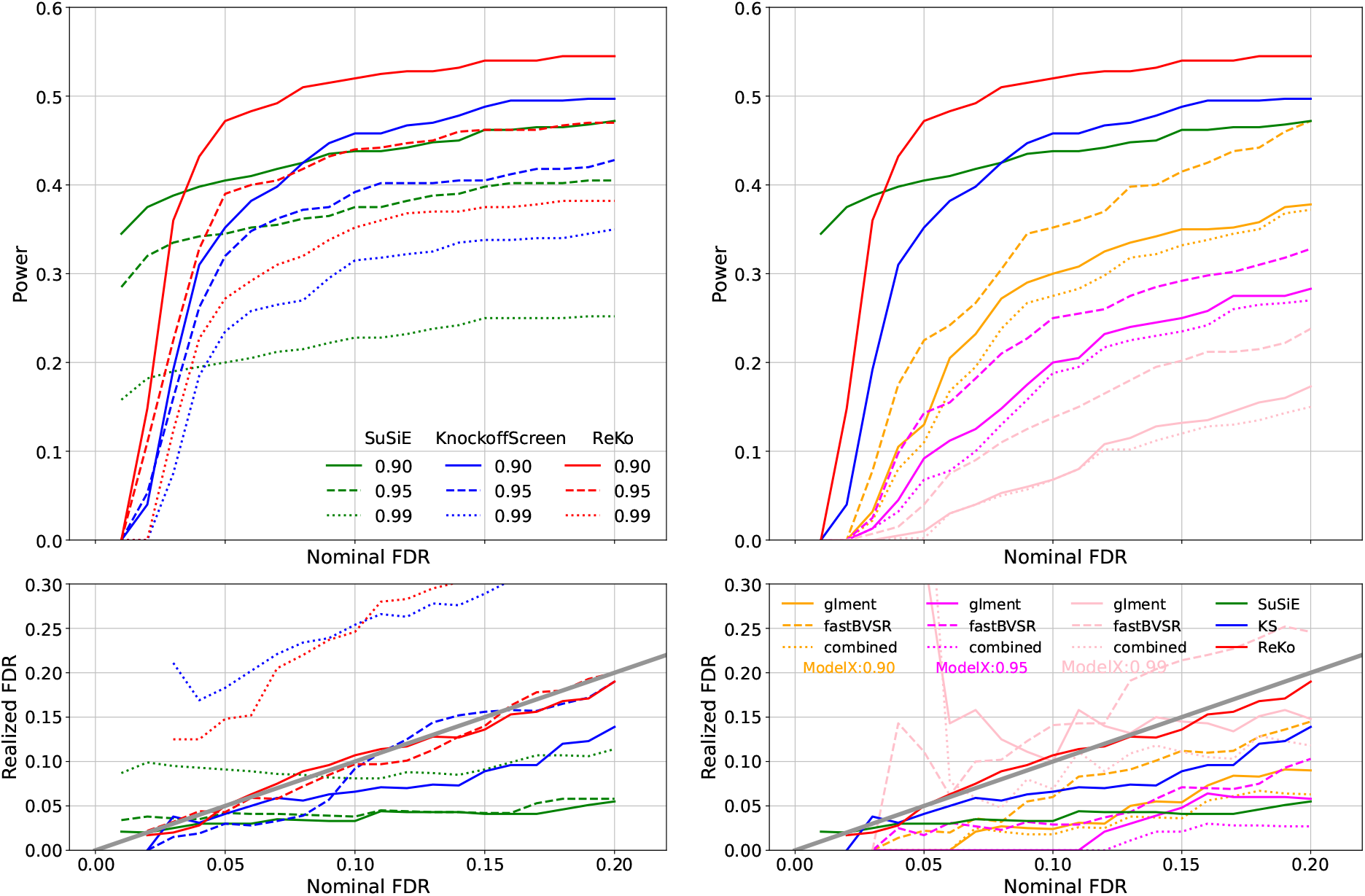
Left panel: Comparisons between SuSiE, KnockoffScreen ⊗combined, and ReKo ⊗combined for different correlation threshold with features were extracted from genomic region *HLA-DQB1*. Results of SuSiE in green; Results of KnockoffScreen in blue; Results of ReKo in red. Different line styles represent thresholds used to trim features. Right panel: Comparison of Model-X ⊗glment (solid line), Model-X fastBVSR (dashed line), and Model-X⊗ combined (dotted line) for different correlation threshold (orange for correlation threshold of 0.90, magenta for 0.95, and pink for 0.99) using the same features extracted from *HLA-DQB1*. Results for SuSiE, KnockoffScreen⊗ combined, and ReKo ⊗combined at the threshold of 0.90 were reproduced on the right panel for comparison. The power and realized FDR were computed from 50 replicates, and each replicate aggregates 10 knockoff statistics.

Figure 3 right panel compares Model-X under different knockoff filters using the same datasets. The first interesting observation is that at correlation 0.99, Model-X shows inflated realized FDR, a phenomenon not seen with proteomic features (Figure 2) and synthetic features (Supplementary Fig. S3 and Fig. S4). The second observation is that Model-X ⊗fastBVSR is more powerful than Model-X⊗glment and ⊗combined, an observation we made with proteomic features (Figure 2). We therefore use knockoff filters Model-X⊗fastBVSR, KnockoffScreen⊗combined, and ReKo⊗combined for real data analysis.

### 2.5 Proteomic association with age

Proteomic signatures of age are an important focus of proteomics and aging studies (Tanaka et al., 2018; Lehallier et al., 2019; Tanaka et al., 2020; Sun et al., 2023). In a recent UK Biobank study using the Olink Explore platform, 1, 944 proteins out of 2, 923 analyzed were significantly associated with age (Sun et al., 2023). Since protein features are highly correlated, a natural question is how many of these features are drivers and how many are hitchhikers. A driver association remains significant when conditioned on other associated features, while a hitchhiker does not. Thus this question can only be addressed by jointly analyzing all features and cannot be answered by single feature tests. Knockoff filter is a framework to select (driver) features while keeping FDR under control. For the knockoff filter to work, however, it is important to construct knockoffs with high power, and the ReKo is designed for this requirement.

Using a subset of proteomic features from UK Biobank (Sun et al., 2023), we investigated how many proteomic features are drivers in their association with age. We first computed the realized kinship matrix for 14, 752 samples based on genotype data using Kindred (Guan and Levy, 2024c). Then we regressed out fixed effects (sex and BMI) and the random effect, whose covariance matrix is a linear scaling of the kinship, from age (the phenotype) and each of the 1, 459 proteomic features using IDUL (Guan and Levy, 2024b). The residuals of age and the proteomic features were used for further analysis. We first examined the marginal association. At a p-value threshold of 3 × 10^−5^, set by Bonferroni correction, there were 902 out of 1, 459 (or 62%) proteomic features significantly associated with age. This proportion aligns with a previous report of 1, 944 out of 2, 923 (or 67%) (Sun et al., 2023).

We applied three knockoff filters to call positives at nominal FDR levels of 0.05 and 0.10. These knockoff filters include Model-X⊗glmnet, KnockoffScreen⊗combined, and ReKo ⊗combined. For each knockoff filter, we aggregated 10 sets of knockoff statistics to call positives (Materials and Methods). (The counts of positives called with a single set of knockoff statistics can be found in Supplementary Tab. S3.) Table 1 summarizes the results. At a nominal FDR of 0.05 Model-X⊗fastBVSR called 256 positives (or 16.9% of total number of features), KnockoffScreen⊗combined called 306 (or 21.0%), ReKo⊗combined called 309 (or 21.2%). At a nominal FDR of 0.10 Model-X⊗fastBVSR called 308 (or 19.9%), KnockoffScree ⊗combined called 361 (or 24.7%), ReKo⊗combined called 371 (or 25.4%).

**Table 1:** Counts of significant association for proteomic data. Column MX contains results from Model-X⊗fastBVSR; Column KS contains results from KnockoffScreen ⊗combined; Column ReKo contains results from ReKo⊗combined. Denote *S*_*K*_ and *S*_*R*_ positive calls by KnockoffScreen⊗combined and ReKo⊗combined respectively, Column Overlap contains counts of intersect between *S*_*K*_ and *S*_*R*_, and the ratios in the parenthesis are Jaccard similarity coefficient.

| Level | MX | KS | ReKo | Overlap |
| --- | --- | --- | --- | --- |
| 0.05 | 256 | 306 | 309 | 291 (90%) |
| 0.10 | 308 | 361 | 371 | 345 (89%) |

At nominal FDR level of 0.05 the number of overlapping positives between KnockoffScreen and ReKo is 291, the number of positives called by either KnockoffScreen or ReKo is 324, and the ratio between counts of intersection and union, known as Jaccard similarity coefficient, is 90%. At nominal FDR level of 0.10 the counts of intersection and union between positives called by KnockoffScreen and ReKo are 345 and 387, and the Jaccard similarity coefficient is 89%. The high Jaccard similarity coefficients signal consistency of positive calls between KnockoffScreen and ReKo. We thus conclude that the majority of proteomic features that show significant association with age in a single feature test are hitchhikers rather than drivers.

### 2.6 Genetic fine mapping

Genome-wide association studies (GWAS) and expression quantitative trait loci (eQTL) analysis are effective at identifying genomic regions associated with phenotypes. Due to linkage disequilibrium, however, pinpointing the exact driver variants within these regions is challenging. Fine mapping methods (Hormozdiari et al., 2014; Benner et al., 2016; Wang et al., 2020) aim to distinguish the driver variants from hitchhiker variants whose association with the phenotype diminishes once conditioned on driver variants. Among current fine mapping methods SuSiE (Wang et al., 2020) is an elegant Bayesian regression approach that models multiple driver variants within genomic regions. Here we demonstrate that knockoff filters can be applied to tackle fine mapping.

In a recent study, we discovered abundant genes harboring eQTL (eGene) that have parent-of-origin effect (POE) (Guan and Levy, 2024a). When a eGene harbors both genotype eQTL and POE eQTL, one may wonder whether POE eQTL are driver signals. Fine mapping can help distinguish drivers from hitchhikers. A challenge in this analysis, however, is that collinearity is introduced into features when genotypes are separated into paternal and maternal alleles. We therefore removed genotypes and only use paternal and maternal alleles in fine mapping. Removing genotypes still allows us to distinguish between POE eQTL and genotype eQTL, because paternal and maternal alleles tend to have similar effects in the absence of POE.

We selected five example eGenes from the POE eQTL study to perform fine mapping. The selections include *PPIEL* and *GJB6* that harbor both genotype eQTL and POE eQTL, *CCR9* that harbors exclusively paternal eQTL, *COA8* that harbors exclusively maternal eQTL, and *NECAB3* that harbors opposing eQTL, whose paternal and maternal alleles having their effect sizes in opposite directions. For each eGene, we regressed out fixed effects including age, sex, BMI, and cell type composition, and random effects that account for relatedness using IDUL (Guan and Levy, 2024b). A consecutive genomic region in cis to the eGenes was visually identified by examining the eQTL peak for each selected eGene. Features in the genomic region were then trimmed such that the largest correlations among the remaining features did not exceed 0.90. If a pair of features displayed a correlation larger than 0.90, we kept the one with stronger marginal association. The trimming was conducted using genotypes, and in our analysis paternal and maternal alleles always present in pairs.

We performed fine mapping with the five selected eGene using four methods: three knock-off filters Model-X⊗fastBVSR, KnockoffScreen⊗combined, and ReKo⊗combined, and SuSiE. Table 2 summarizes the results and from which we make the following observations: 1) *PPIEL* is known to have both genotype eQTL and paternal eQTL. All four methods identified two paternal alleles as drivers. 2) *GJB6* is also known to have both genotype eQTL and maternal eQTL. Both paternal and maternal alleles of SNP *rs12875121* were found to be significant by all three knockoff methods. SuSiE missed the positive call of maternal allele by 0.01. This SNP is indeed a genotype eQTL. 3) ReKo called two additional maternal eQTL significant for *GJB6*, but Model-X, KnockoffScreen, and SuSiE did not. Regression analysis show that paternal and maternal alleles of SNP *rs12875121* alone had adjusted R-squared value of 0.081, and adding these two maternal eQTL the adjusted R-squared value increased to 0.103. This suggests that the two maternal maternal eQTL are likely true positives. 4) *CCR9* is known to harbor exclusively paternal eQTL, and all four methods call a paternal eQTL positive. 5) *COA8* provides another example where all three knockoff filters called positive, but the SuSiE missed by a small margin. 6) *NECAB* is known to harbor exclusively opposing eQTL; all four methods reported significant maternal eQTL without its companion paternal eQTL.

**Table 2:** Fine mapping of selected eGenes. MX: Model-X; KS: KnockoffScreen. Column eQTL contain variants information in the format of rs:ref:alt:chr:pos:pm where pm can be 1 for paternal alleles, and 0 for maternal alleles. Columns MX, KS, and ReKo contain average knockoff statistics from knockoff filter ⊗fastBVSR, aggregated from 10 replicates of knockoffs respectively. Column SuSiE contains PIP from SuSiE. Column Type contain four binary digits with 1 for positive call and 0 for negative call. From left to right the first digit is for Model-X ⊗fastBVSR, the second digit is for KnockoffScreen ⊗combined, the third is for ReKo⊗ combined, and the fourth digit is for SuSiE. The nominal FDR used to call positives is 0.10.

| eGene | eQTL | MX | KS | ReKo | SuSiE | Type |
| --- | --- | --- | --- | --- | --- | --- |
| *PPIEL* | rs79473113:C:A:1:39554348:1 | 1.00 | 1.00 | 1.00 | 1.00 | 1111 |
|  | rs7520271:A:C:1:39579041:1 | 0.80 | 0.82 | 0.87 | 0.97 | 1111 |
| *GJB6* | rs12875121:T:C:13:20478719:1 | 0.95 | 0.97 | 0.98 | 0.99 | 1111 |
|  | rs12875121:T:C:13:20478719:0 | 0.76 | 0.78 | 0.79 | 0.89 | 1110 |
|  | rs9506450:G:C:13:20239478:0 | -0.15 | -0.27 | 0.61 | 0.44 | 0010 |
|  | rs9579911:C:G:13:20637437:0 | 0.23 | 0.26 | 0.38 | 0.20 | 0010 |
| *CCR9* | rs9814578:T:G:3:45821775:1 | 0.85 | 0.93 | 0.98 | 0.98 | 1111 |
| *COA8* | rs10143623:G:A:14:103720539:0 | 0.66 | 0.58 | 0.85 | 0.87 | 1110 |
| *NECAB3* | rs2626556:A:G:20:33768509:0 | 0.79 | 0.80 | 0.88 | 0.95 | 1111 |

## 3 Discussion

In this paper, we presented a novel method to construct knockoffs via Householder reflection and demonstrated the power it brings to knockoff filters. We provided a simple method to aggregate multiple sets of identically distributed knockoff statistics to improve the consistency of knockoff filters. We also demonstrated that a combined knockoff filter, by combining Bayesian and Frequentist inference, can effectively bring the realized FDR under control when correlation is not extremely high. ReKo assumes feature matrix is fixed and known. In this sense, ReKo works within the framework of Fixed-X but provides a novel way to estimate **S** in Equation 1. Our current work, however, provides no theoretical guarantee of FDR control under arbitrary covariance structure.

We applied reflection knockoffs to address two problems that involve highly correlated features. In the analysis of proteomic signatures of age using knockoff filters, we demonstrated that the majority of proteins showing significance in single-feature tests are hitchhikers, not drivers. In the analysis of genetic fine mapping, we demonstrated that ReKo can reliably make positive calls that are consistent with Model-X and KnockoffScreen. Two extra positive calls by ReKo alone were supported by high SuSiE PIPs. ReKo is computationally efficient, particularly when rSVD approximation is invoked. We documented the computational time required to generate knockoffs and to fit models for different knockoff filters in Supplementary Section 10.

In both simulated features and real datasets, ReKo improves the power of different knock-off filters. We believe ReKo can be beneficial to other knockoff-based methods, such as those developed for genetic association studies (Candés et al., 2018; He et al., 2021b). In Figure 1, we demonstrate that the aggregation helps KnockoffScreen (He et al., 2021a). We believe our method of aggregating multiple sets of identically distributed knockoff statistics can also benefit knockoffs constructed by other methods, including hidden Markov model knockoffs (Sesia et al., 2019), KnockoffZoom (Sesia et al., 2020), and GhostKnockoffs (He et al., 2022).

We used empirical Bayes estimated shrinkage priors to combat high correlation when constructing reflection knockoffs, an alternative approach is to use weighted regression in Equation (9) to down-weight covariates in **X**_−*j*_ that are highly correlated with **X**_*j*_. This is equivalent to putting a different prior on *β*_−*j*_ for different *j*. A natural prior for this purpose would be the partial correlation taken from cov(**X**)^−1^. We would still need a separate shrinkage prior to control the overall shrinkage (similar to the *λ* in LASSO), which could be determined empirically, for example, using cross-validation. This can be a fruitful future research topic to further improve reflection knockoffs.

Trimming highly correlated features was used in the current study to reduce high correlation in the data. As currently implemented, the trimming procedure aimed to keep features with more significant marginal association. There is certainly room for improvement, such as fitting models jointly to rank features, and using the ranks to inform feature trimming. Compared to trimming, it may be more fruitful to cluster highly correlated features or to perturb highly correlated features with carefully designed noises. This can be another fruitful future research topic to improve reflection knockoffs.

A clever approach to reduce high correlation in fine mapping is to pool genetic data from multiple ancestral backgrounds (Yuan et al., 2024; Cai et al., 2023). This practice takes advantage of different patterns of LD in different ancestral backgrounds to reduce correlation among genetic markers, which evidently increases power for fine mapping. Knockoff filters can also benefit from pooling genetic data from multiple ancestral backgrounds.

A Bayesian procedure such as fastBVSR may produce PIPs that are not calibrated due to, say, difficulties in estimating priors odds in applications involving highly correlated features. Our study suggests that knockoff filters can indeed improve calibration of PIP by recalibrating it. Similar to computing p-values for Bayes factors (Zhou and Guan, 2018), using a knockoff filter – an intrinsic Frequentist procedure – with PIP as the importance measure can be perceived as another example of Bayesian / non-Bayesian compromise (Good, 1992). Another notable benefit of the knockoff filter is its ability to combine multiple knockoff filters, such as ⊗fastBVSR and ⊗glmnet, to improve calibration of the realized FDR. The combined knockoff filter can also be perceived as an example of Bayesian / non-Bayesian compromise.

Both knockoff filters ReKo⊗glmnet⊗Beta and ReKo⊗fastBVSR⊗PIP produced inflated realized FDR when features were highly correlated. The combined knockoff filter appeared to be effective in controlling the realized FDR when the correlation is not extremely high. The intuition behind this is as follows: Suppose there are four highly correlated features such that conditioning on any one of these features, the other three features become uncorrelated with the phenotype. One of the features is selected as a driver in simulation. Penalized regression will arbitrarily pick one feature among the four and call it a positive, while the other three are declared negative. In contrast, fastBVSR, by the virtue of model averaging, will assign each feature a PIP of about 0.25. If the knockoff threshold happens to be 0.20, then fastBVSR will produce three false positives, compare to at most one for penalized regression. On the other hand, suppose these four features are null features, only associated with the phenotype via hitchhiking. The model averaging of fastBVSR will make it less likely to produce false positives among these four features compared to glmnet. The combined filter will reduce false positive in both scenarios.

One potential application reflection knockoffs is data sharing for genetic research. By design, knockoffs ensure that the original genetic data cannot be reverse-engineered, thanks to their reliance on random processes and the inherently one-way nature of the construction method. Even with complete knowledge of the algorithm and access to the knockoff data, recovering the original data is mathematically infeasible. This opens a transformative pathway for sharing sensitive genetic or proteomic data while meeting stringent privacy and regulatory standards, such as HIPAA. Imagine a collaborative research ecosystem where scientists worldwide can freely access anonymized knockoff datasets to address critical challenges like fine mapping. This mirrors the ethos of the machine learning field, where shared data fuels innovation: the data are there, the goals are clear – innovate with dare! With reflection knockoffs, genetic research can embrace this model, empowering the community to collectively tackle the most pressing problems with creativity and bravery.

## 4 Materials and Methods

### 4.1 Reflection

We have the following lemma regarding Householder reflection (c.f. Golub and Van Loan, 2013) and some useful properties for constructing knockoffs.

#### Lemma 1.

*Suppose the data matrix* **X** *is n* × *p with n* > *p and* **X** *is column-centered, scaled, and has full rank p. Denote* 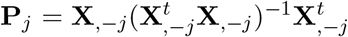 *where* **X**_,−*j*_ *is matrix* **X** *with j-th column removed. Compute* **Y** *such that its j-th column is a Householder reflection of* **X**_*j*_ *with respect to* **P**_*j*_, **Y**_*j*_ = (2**P**_*j*_ − **I**)**X**_*j*_. *Define* 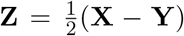. *Denote cov*(**X**_*j*_, **Y**_*k*_) *as sample covariance between vector* **X**_*j*_ *and* **Y**_*k*_. *Then the following hold:*

1. *cov*(**Y**_*j*_, **X**_*k*_) = *cov*(**X**_*j*_, **X**_*k*_) *for all j* ≠*k and j, k ∈* (1, 2, …, *p*).
2. *cov*(**Y**_*j*_, **X**_*j*_) < *var*(**X**_*j*_) *for all j ∈* (1, 2, …, *p*).
3. *Let* **T** *be a matrix such that* **T**_*jk*_ = *cov*(**X**_*j*_, **Z**_*k*_), *then* **T**_*jk*_ = 0 *for all j* ≠ *k and* **T**_*jj*_ = *cov*(**Z**_*j*_, **Z**_*j*_) > 0 *for all j*.
4. **X**(**X**^*t*^**X**)^−1^**XY** = **Y** *and* **X**(**X**^*t*^**X**)^−1^**XZ** = **Z**.

*Proof*. a) The projection matrix 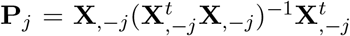 projects **X**_*j*_ onto the column space of **X**_,−*j*_. Since (**I**−**P**_*j*_)**X**_*j*_ is orthogonal to **X**_*k*_ for *k* ≠ *j*, we have cov((**I**−**P**_*j*_)**X**_*j*_, **X**_*k*_) = 0. This implies cov(**P**_*j*_**X**_*j*_, **X**_*k*_) = cov(**X**_*j*_, **X**_*k*_). Now **Y**_*j*_ = (2**P**_*j*_ − **I**)**X**_*j*_, and direct computation gives: cov(**Y**_*j*_, **X**_*k*_) = cov((2**P**_*j*_ − **I**)**X**_*j*_, **X**_*k*_) = 2cov(**P**_*j*_**X**_*j*_, **X**_*k*_) − cov(**X**_*j*_, **X**_*k*_). Substituting cov(**P**_*j*_**X**_*j*_, **X**_*k*_) = cov(**X**_*j*_, **X**_*k*_), we obtain cov(**Y**_*j*_, **X**_*k*_) = cov(**X**_*j*_, **X**_*k*_), proving (a).

b) The projection of **X**_*j*_ onto the space spanned by **X**_,−*j*_ as **X**_**V**_ = **P**_*j*_**X**_*j*_, and the or-thogonal component as **X**_*⊥*_ = (**I** − **P**_*j*_)**X**_*j*_. Thus, **X**_*j*_ = **X**_**V**_ + **X**_*⊥*_. The Householder reflection **Y**_*j*_ can then be written as **Y**_*j*_ = **X**_**V**_ − **X**_*⊥*_. The sample covariance between **Y**_*j*_ and **X**_*j*_ is given by cov(**Y**_*j*_, **X**_*j*_) = cov(**X**_**V**_ − **X**_*⊥*_, **X**_**V**_ + **X**_*⊥*_) = cov(**X**_**V**_, **X**_**V**_) − cov(**X**_*⊥*_, **X**_*⊥*_). That is, cov(**Y**_*j*_, **X**_*j*_) = var(**X**_**V**_) − var(**X**_*⊥*_). Since var(**X**_*j*_) = var(**X**_**V**_) + var(**X**_*⊥*_), it follows that: cov(**Y**_*j*_, **X**_*j*_) = var(**X**_*j*_) − 2var(**X**_*⊥*_). Since var(**X**_*⊥*_) *≥* 0 with equality holds only if **X**_*⊥*_ = 0, which would violate the assumption that matrix **X** has full rank. Hence, cov(**Y**_*j*_, **X**_*j*_) < var(**X**_*j*_), proving (b).

c) Statement c) follows directly from a) and b). Specifically, for any *j*, 4var(**Z**_*j*_) = cov(**X**_*j*_ − **Y**_*j*_, **X**_*j*_ − **Y**_*j*_) = cov(**X**_*j*_)+cov(**Y**_*j*_)− 2cov(**X**_*j*_, **Y**_*j*_) = 2− 2cov(**X**_*j*_, **X**_*j*_ − 2**Z**_*j*_) = 4cov(**X**_*j*_, **Z**_*j*_) = 4**T**_*jj*_. From (b) cov(**X**_*j*_, **Y**_*j*_) < var(**X**_*j*_), ensuring **T** = cov(**X, Z**) > 0.

d) From (a), **Y**_*j*_ = (2**P**_*j*_−**I**)**X**_*j*_, where **P**_*j*_ projects **X**_*j*_ onto the column space of **X**_,−*j*_. Thus, **Y**_*j*_ is a linear combination of columns of **X**. Consequently, **Y** lies in the column space of **X**. Similarly, 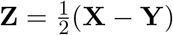 also lies in the column space of **X**. Therefore: **X**(**X**^*t*^**X**)^−1^**XY** = **Y** and **X**(**X**^*t*^**X**)^−1^**XZ** = **Z**.

### 4.2 Construct knockoffs based on reflection

Following Equation (2.2) of (Barber and Candés, 2015) and Section 3.1.1 of (Candés et al., 2018), the mean **b** and covariance **C**^*t*^**C** of a knockoff 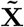 conditional on **X** are given by

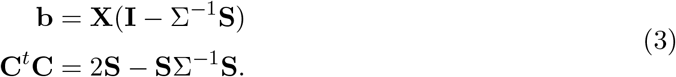

Let **S** = *α***T** for *α ∈* (0, 1]. Straightforward calculations using the identities in Lemma 1 parts (c) and (d) yields:

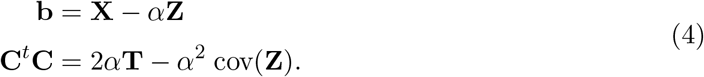

To ensure **C**^*t*^**C** *≥* **0**, we need to determine 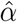 such that this condition holds. Noting that **T** is positive definite and diagonal, we can rewrite:

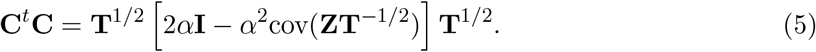

Let **Q**Λ**Q**^*t*^ be the eigendecomposition of cov(**ZT**^−1*/*2^). Substituting this, we obtain

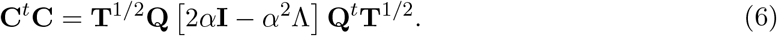

For **C**^*t*^**C** to be positive semi-definite, it is sufficient to have

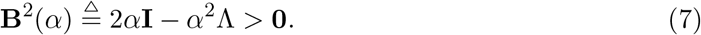

Therefore it’s sufficient to have *α ≤* 2*/*Λ_1_, where Λ_1_ is the leading eigenvalue of cov(**ZT**^−1*/*2^). Since Λ_1_ is positive and finite, and *α ∈* (0, 1], the optimal 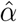 exists. This ensures **C**^*t*^**C** is positive semi-definite, with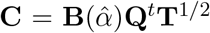. Consequently, 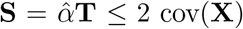, as guaranteed by the Schur complement condition.

We thus construct the knockoff as

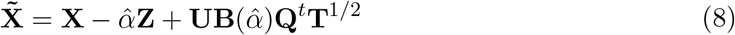

where: If **U** is chosen from the null space of 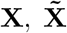 is akin to Fixed-X knockoffs. If 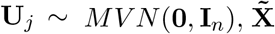 is akin to Model-X second-order knockoff. Our numerical studies showed that their performances are highly similar (Supplementary Tab. S2), and in this paper we use the later for numerical convenience. As a special case, when the columns of **X** are uncorrelated, simple calculations show that 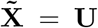. This implies that for uncorrelated 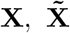 can be simulated directly from either a standard normal distribution or the basis of the null space of **X**.

### 4.3 Shrinkage priors and data augmentation

In Lemma 1, computing **Y**_*j*_ requires the least square fit of the linear regression:

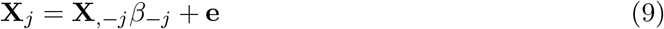

to obtain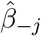, which is then used to calculate 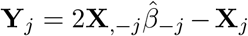. When *p* is large compare to *n*, and/or there exist highly correlated features, the model (9) is prone to overfitting, which can adversely affect the performance of the knockoff filter. While penalized regression is a good choice to avoid overfitting, Bayesian regression offer a more elegant solution in this context, as it serves two purposes.

Suppse we put priors on *β*_−*j*_ such as

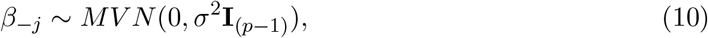

where *MV N* denotes multivariate normal distribution, **I**_*m*_ is the identity matrix of dimension *m*, and *σ* is the prior effect size to be specified. The Bayesian estimated effect sizes are given by

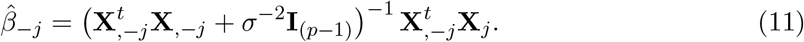

The same solution can be achieved via augmenting **X** with ghost observations *σ*^−1^**I**_*p*_ such that **X** *←* [**X**^*t*^, *σ*^−1^**I**]^*t*^. A natural choice for *σ* is *σ*^2^ = 1*/p*, as this prior ensures the posterior distribution remains well behaved for large *p*. That is, the posterior remain proper and without excessive variance inflation. But empirical Bayes estimated *σ* (to be discussed below) appears to perform better. This data augmentation enables constructing reflection knockoffs for **X** of any shape, as the augmented **X** will have dimension (*n* + *p*) × *p* and full rank *p*, which satisfies the conditions in Lemma 1. By augmenting **X** with ghost observations, we can extend this construction to handle cases where *p* > *n*.

### 4.4 Empirical Bayes estimates of shrinkage priors

Consider a Bayesian linear regression for model (9) and prior (10), we have

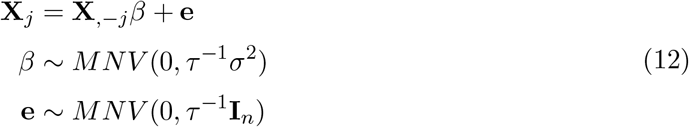

Integrate out *β* to obtain the marginal likelihood, and we get

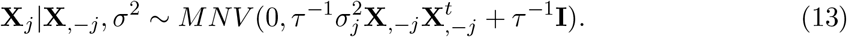

This is a special case of a linear mixed model and we can obtain maximum likelihood estimate of 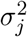 using IDUL (Guan and Levy, 2024b). To proceed, we obtain eigeindecomposition 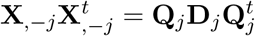 where **Q**_*j*_ is orthonormal and **D**_*j*_ is diagonal. We rotate the system by multiplying (12) by **Q**^*t*^ from left we get

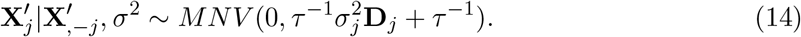

For this particular application, we find that using a bisection method is more stable than IDUL, particularly for smaller *p*. In addition, a practical upper bound 1 can be imposed on 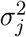 to facilitate the bisection. Therefore, we used the bisection method to find maximum likelihood estimate of *σ*_*j*_.

We estimated 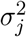 for each feature, which requires eigendecomposition of 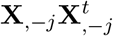 for each *j*, and naive computation can be expensive *O*(*np*^3^). Here we approximate eigenvectors of 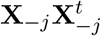 with eigenvectors of **XX**^*t*^, but update eigenvalues for each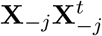. The rank one update for eigenvalue requires one to solve secular equations, which can be solved in a constant time (for updating a single eigenvalue) using another bisection search method. The total complexity of to obtain eigenvalues for 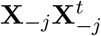 using rank one update is *O*(*np* + *p*^2^) (more details in Supplementary Section 1). For eigen decomposition of **XX**^*t*^, we implemented randomized singular value decomposition (rSVD) (Halko et al., 2011) to efficiently compute (say 200) approximate leading eigenpairs, and use them to compute empirical Bayes estimated priors.

After estimating *σ*_*j*_ for each *j*, we obtained Σ = *diag*(*σ*_1_, …, *σ*_*p*_). Instead of fitting Model (12) for each *j*, which would be prohibitively, we augmented **X** with Σ and compute reflection using Algorithm 1 detailed below. This appears to maintain the performance and computation efficiency. A comparison between data augmentation with empirical Bayes estimated priors and those with fixed priors can be found in Supplementary Fig. S1.

### 4.5 Computation

To speed up the computation and avoid repeatedly computing the matrix inverse needed for 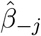 in Equation (11), we employ QR decomposition of **X**: **X** = **QR**, where **Q** is orthonormal *n* × *p* and **R** is a *p* × *p* upper triangular matrix. Multiply both sides of the linear regression (9) by *Q*^*t*^, we get **R**_*j*_ = **R**_,−*j*_*β*_−*j*_ + **e**_**Q**_, where **R**_*j*_ is the *j*th column of **R**, and **R**_,−*j*_ is the matrix with its *j*-th column removed.

#### Efficient computation via Givens Rotation

The matrix **R**_,−*j*_ is close to being upper triangular and requires at most (*p* − *j*) many Givens rotations to become triangular, which can be done efficiently. A Givens rotation corresponds to left-multiplying a sparse matrix, which we apply to both sides of the regression. Importantly, this rotation is unitary and does not affect mean and variance of the error term. Once the matrix is properly triangularized, we use backward substitution to solve the system for 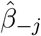. This value is then used to compute 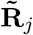 and subsequently **Y**_*j*_.

#### Parallel Computing for Speed-Up

The algorithm below outlines the steps, where the for loop can be parallelized for efficiency. Our software implementation leverages this feature for high-performance computation.

##### Algorithm 1

QR decomposition for Householder reflection

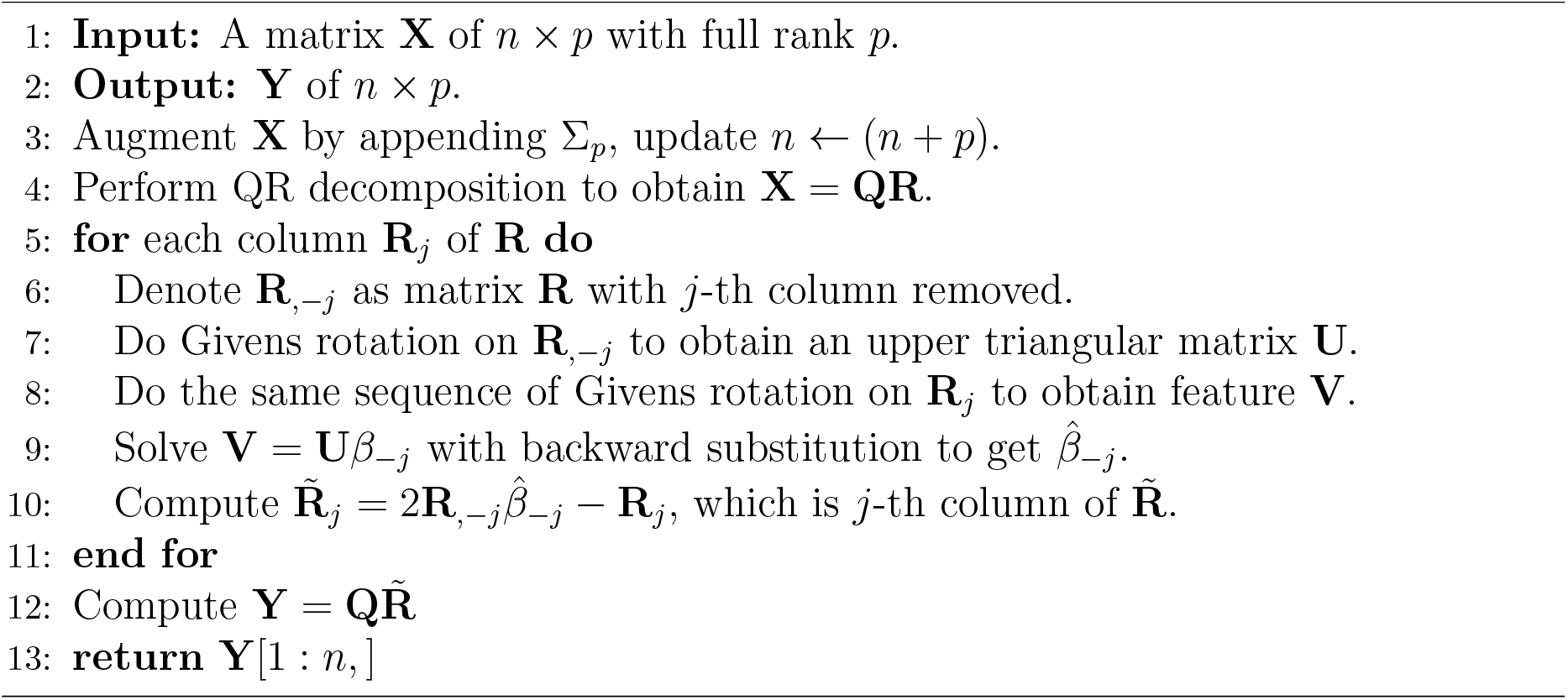

### 4.6 Aggregation

We present a method for aggregating multiple knockoff statistics. Let *W* = (*W*_1_, …, *W*_*p*_) represent the knockoff features importance statistics from a single reflection knockoff. Under the null, the distribution *W*_*j*_ is symmetric about zero, ensuring FDR control. The knockoff threshold *T* for *W*, controlling the FDR at nominal level *q* is defined as:

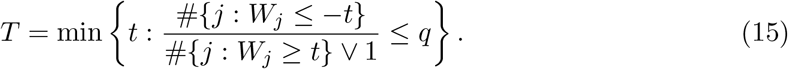

Suppose we construct *M* copies of reflection knockoffs, the *m*-th copy of the knockoff lead to the knockoff statistics *W* ^(*m*)^ for *m* = 1, …, *M*, which are identically distributed as *W*. This assumes the same knockoff construction mechanism, data, and model-fitting procedure, differing only by random seeds or knockoff randomness. The following lemma establishes the validity of aggregating these identically distributed knockoff statistics.

#### Lemma 2.

*Let W*_*cat*_ = (*W* ^(1)^, …, *W* ^(*M*)^), *i*.*e*., *the pooled knockoff statistics from the M runs. If we apply the standard knockoff threshold selection rule* (15) *to W*_*cat*_, *the resulting threshold T*_*cat*_ *also controls the FDR at level q. Define the average knockoff statistic* 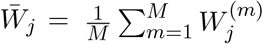. *Selecting feature j such that* 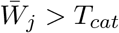 *controls the FDR at level q*.

*Proof*. For the *m*-th knockoff statistics *W* ^(*m*)^, the null features are symmetrically distributed around zero and the knockoff threshold selection 15 controls the FDR at level *q* for *W* ^(*m*)^. Since *W* ^(*m*)^ for *m* = (1, …, *M*) are identically distributed, the combined set *W*_*cat*_ of size *Mp* is drawn from the same underlying distribution as *W*. If the *j*-th feature is a null feature, then 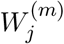 is symmetric about zero for all *m ∈* (1, …, *M*). Conversely, if the *j*-th features is non-null, then 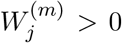 for all *m ∈* (1, …, *M*). The two counts in (15) have the same expectation for each *j* and for all *t*. Thus, applying the knockoff threshold rule to *W*_*cat*_ yields a threshold *T*_*cat*_ that controls FDR at level *q*.

For a null feature *j*, the distribution of 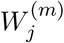 is symmetric about zero. Formally, the pair 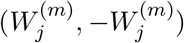 is identically distributed. This symmetry holds for each copy of the knockoff. The average statistics 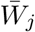 preserves this symmetry, ensuring that 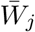 is symmetry about zero. The super-martingale argument from the original FDR control proof applies directly (supplementary material of Barber and Candés, 2015). Under the null, features have no preference for positive or negative deviations. Thus, selecting feature *j* such that 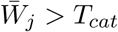 controls the FDR at level *q*.

## 5 Acknowledgements

This research is supported by the Division of Intramural Research of the National Heart, Lung, and Blood Institute, Bethesda, MD (D.L. Principal Investigator). We thank editors and two anonymous reviewers for their constructive review comments that greatly improved clarity of the presentation. In particular, we thank one reviewer who brought our attention to KnockoffScreen, and the other for suggesting empirical Bayes estimates of the prior effects for reflection. Our revised manuscript benefitted greatly from both suggestions.

## 7 Data and code availability

This study analyzed existing datasets from public domain and simulated datasets. Data from the UK Biobank can be obtained from https://www.ukbiobank.ac.uk/, and datasets from Framingham Heart Study can be obtained via dbGaP. The software for ReKo and its source code are freely available at https://github.com/haplotype/ReKo.

## 8 Author contributions

Y.G. conceived the study, developed the methodology, implemented the computational software and performed experiments, analyzed results, and wrote the manuscript. D.L. supervised the work, provided access to data and computation, edited and approved the final manuscript.

