## Supplementary for "Reflection Knockoffs via Householder Reflection: Applications in Proteomics and Genetic Fine Mapping"

May 27, 2025

### 1 Rank-one Update for Eigenvalues and Eigenvectors

For ReKo we need to obtain eigendecomposition of $\mathbf{X}_{-j}\mathbf{X}_{-j}^t$ for each $j$, and naive computation can be expensive $O(np^3)$. Notice that $\mathbf{X}_{-j}\mathbf{X}_{-j}^t = \mathbf{X}\mathbf{X}^t - \mathbf{X}_j\mathbf{X}_j^t$, that is a rank-one subtraction of $\mathbf{X}\mathbf{X}^t$. We can use reverse rank-one update to obtain eigendecomposition of $\mathbf{X}_{-j}\mathbf{X}_{-j}^t$ with additional complexity of $O(p)$ from eigendecomposition of $\mathbf{X}\mathbf{X}^t$. This makes the total complexity of $O(np^2+p^2)$. To update new eigenvalues, we want to solve

$$\begin{aligned}
\det(\mathbf{X}_{-j}\mathbf{X}_{-j}^t - \lambda\mathbf{I}) &= \det(\mathbf{X}\mathbf{X}^t - \mathbf{X}_j\mathbf{X}_j^t - \lambda\mathbf{I}) \\
&= \det(\mathbf{Q}(\mathbf{D} - \lambda\mathbf{I})\mathbf{Q}^t - \mathbf{X}_j\mathbf{X}_j^t) \\
&= \det((\mathbf{D} - \lambda\mathbf{I}) - \mathbf{Q}^t\mathbf{X}_j\mathbf{X}_j^t\mathbf{Q}) \\
&= \det((\mathbf{D} - \lambda\mathbf{I}) - vv^t) \\
&= \det(\mathbf{D} - \lambda\mathbf{I})(1 - v^t(\mathbf{D} - \lambda\mathbf{I})^{-1}v) \\
&= 0,
\end{aligned} \tag{1}$$

where $v = \mathbf{Q}^t\mathbf{X}_j$. To this end, we want to solve secular equation

$$f(\lambda) \triangleq 1 - \sum_{j=1}^{p} \frac{v_j^2}{\mathbf{D}_j - \lambda} = 0 \tag{2}$$

for $\lambda$ where $\mathbf{D}_j$ is the $j$-th diagonal element of $\mathbf{D}$. Following the convention, assume diagonal entries of $\mathbf{D}$ is in decreasing order. These derivations are standard [c.f. 1]. At interval $x \in (\mathbf{D}_{j+1}, \mathbf{D}_j)$, $f(x)$ is continuous, with $f(\mathbf{D}_{j+1} + \epsilon)$ and $f(\mathbf{D}_{j+1} - \epsilon)$ having opposite sign for some small positive $\epsilon$, so that there exists a solution such that $f(x) = 0$. On interval $x > \mathbf{D}_1$ no solution exits, but on interval $x \in [0, \mathbf{D}_p)$ solution exists. On each interval, golden ratio bisection algorithm can be used to solve for an eigenvalue.

Once an updated eigenvalues $\lambda'$ is obtained, we can update eigenvectors. From $\mathbf{X}_j = \mathbf{Q}v$, then eigenvalue system becomes $\mathbf{Q}(\mathbf{D} - vv^t)\mathbf{Q}^t u' = \lambda' u'$. Define $w = \mathbf{Q}^t u'$, thus $u' = \mathbf{Q}w$. Then eigenvalue system becomes $(\mathbf{D} - vv^t)w = \lambda' w$. Rearrange to get $(\mathbf{D} - \lambda')w = v(v^t w)$. Denote $a = v^t w$, we get

$$w = (\mathbf{D} - \lambda')^{-1} v. \tag{3}$$

We compute $u' = \mathbf{Q}w$ and renormalize it to have unit variance.

#### 2 Simulation Correlated Features

We designed two schemes to simulate correlated features. One is equi-correlation $c$ between all pairs of features, where we examined $c = 0.2, 0.5, 0.8$ representing low, medium, and high levels of correlation. The other is vary-correlation where each pair of features have their correlations drawn from a bell-shaped histogram, and we let histogram peak at $0.2, 0.5, 0.8$. Vary-correlation better mimics the real data, but the equal correlation were used in the seminal paper of knockoff [2], and we include here for comparison (Tab. S2). The weights for histograms peaked at $0.2, 0.5$ and $0.8$ are shown in Table S1 below.

| Correlation | 0 | 0.1 | 0.2 | 0.3 | 0.4 | 0.5 | 0.6 | 0.7 | 0.8 | 0.9 |
|---|---|---|---|---|---|---|---|---|---|---|
| Low | 20 | 40 | 60 | 40 | 20 | 10 | 0 | 0 | 0 | 0 |
| Medium | 0 | 0 | 10 | 20 | 40 | 60 | 40 | 20 | 10 | 0 |
| High | 0 | 0 | 0 | 0 | 0 | 10 | 20 | 40 | 60 | 40 |

Tab. S1: Weights used to draw correlation between samples. Low correlation has range $(0.1, 0.6)$ with peak at 0.2, Median has range $(0.2, 0.8)$ with peak at 0.5, and High has range $(0.5, 0.9)$ with peak at 0.8.

In addition to simulate multivariate normal distributed features, we also simulated multivariate $t$ distributed features with degree of freedom 5. This is done by scaling the multivariate normal vectors by a factor that is drawn from a $t$ (d.f.5) distributed random variable.

For sample size $n = 2000$ and number of features $p = 800$ (or $p = 400$), we randomly pick 20 driver features. Their effect sizes were assigned as 1, but then rescaled so that the percentage of phenotypic variation explained (PVE) by these driver features are at a prescribed level. We examined both low (0.20) and high (0.8) PVEs. For each setting of simulation parameters, we generated 100 replicates of features and their associated phenotypes. Different knockoff methods were then used to generate their corresponding knockoffs. Original features and their knockoffs were fed to knockoff filters $\otimes$glmnet to generate knockoff statistics. These 100 sets of knockoff statistics were pooled to call positives at different nominal FDR levels. The called positives were compared with truth to obtain true and false positives.

### 3 Power and Realized FDR with Different Priors

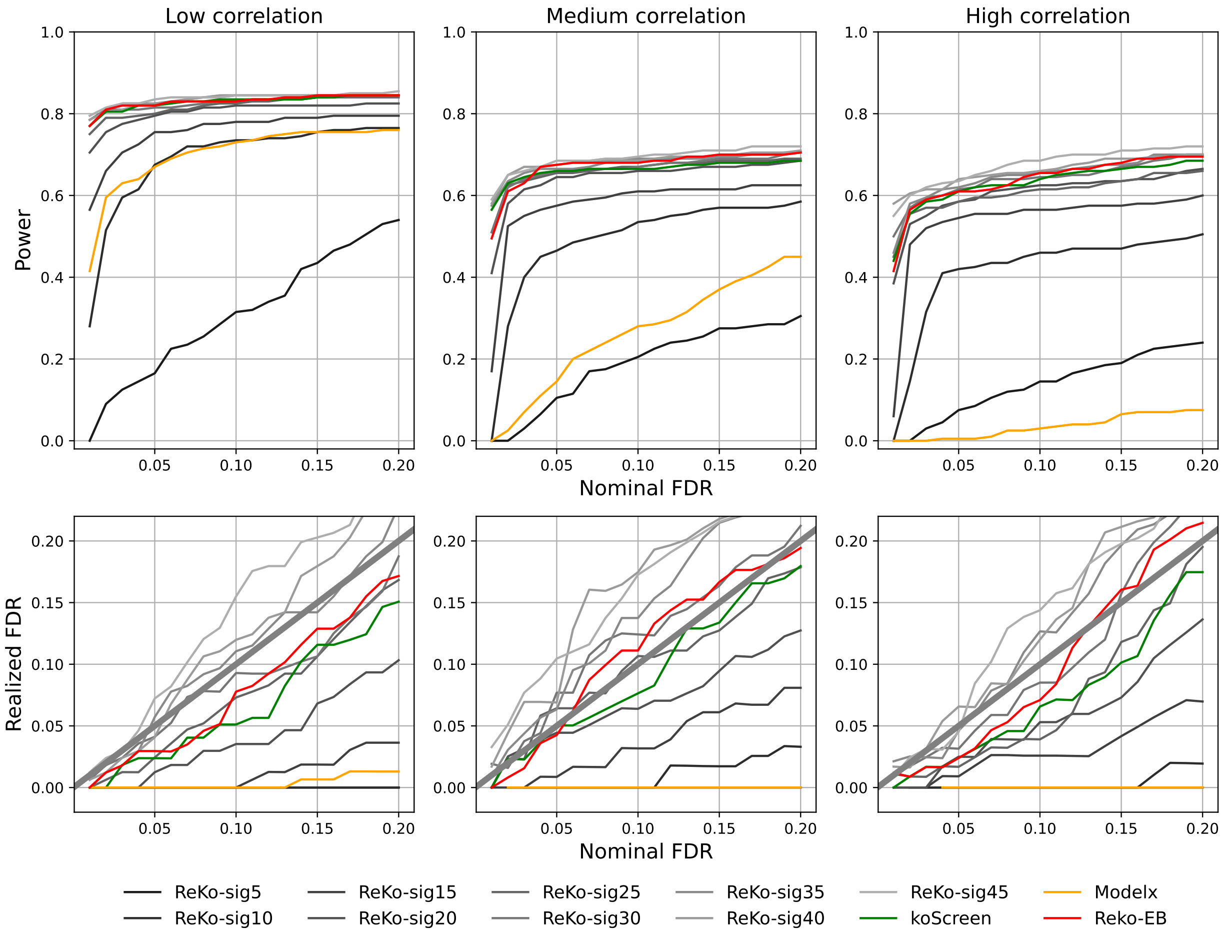


Fig. S1: Power and realized FDR for ReKo under different priors. Model-X (orange), KnockoffScreen (green), and ReK-EB with emprical Bayes-estimated priors (red) are included for comparison. The prior for ReKo reflection is $\sigma \mathbf{I}_p$, and In the legend ReKo-sig5 means $\sigma = 5$. For this simulation $n = 2000$ and $p = 800$ with medium correlation, and $\sqrt{p} \approx 28$. We can see that the prior $\sqrt{p}\,\mathbf{I}_p$ is a sensible choice, but perhaps less powerful than EB-estimated prior.

### 4 Comparison with Fixed-X and Model-X knockoffs

Table S2 compares different knockoffs using knockoff filters ⊗glmnet. For the equi-correlation (the top half of the table), where all pairs of features have identical correlation, the power gain of reflection knockoffs (ReKo-Null and ReKo-Norm) over Fixed-X and Model-X knockoffs was less than in the vary-correlation (the bottom half of the table). Presumably, the semidefinite program used by both Fixed-X and Model-X knockoffs performs better under the equi-correlation scenario, where there are $p-1$ identical small eigenvalues and one large eigenvalue for $\mathrm{cov}(\mathbf{X})$. For varying correlation, ReKo-Null⊗glmnet and ReKo-Norm⊗glmnet produced higher power than both Fixed-X⊗glmnet and Model-X⊗glmnet. In particular, when correlations among features are large, both Fixed-X⊗glmnet and Model-X⊗glmnet have little to no power, while ReKo-Null⊗glmnet and ReKo-Norm⊗glmnet maintain considerable power.

| Mode | Corr | Level | Fixed-X | Model-X | ReKo-Null | ReKo-Norm |
|---|---|---|---|---|---|---|
| Equi-Corr | 0.2 | 0.05 | 0.76 (0.04) | 0.76(0.04) | 0.77 (0.04) | 0.77 (0.06) |
| | | 0.10 | 0.78 (0.10) | 0.78(0.09) | 0.78 (0.09) | 0.79 (0.10) |
| | 0.5 | 0.05 | 0.58 (0.06) | 0.57(0.05) | 0.57 (0.04) | 0.58 (0.06) |
| | | 0.10 | 0.61 (0.13) | 0.60(0.10) | 0.60 (0.11) | 0.62 (0.13) |
| | 0.8 | 0.05 | 0.28 (0.04) | 0.26 (0.04) | 0.31 (0.08) | 0.31 (0.07) |
| | | 0.10 | 0.32 (0.10) | 0.31 (0.07) | 0.34 (0.12) | 0.35 (0.13) |
| Vary-Corr | Low | 0.05 | 0.00 (0.00) | 0.58 (0.00) | 0.76 (0.04) | 0.76 (0.04) |
| | | 0.10 | 0.00 (0.00) | 0.67 (0.03) | 0.79 (0.09) | 0.79 (0.08) |
| | Medium | 0.05 | 0.00 (0.00) | 0.13 (0.00) | 0.67 (0.05) | 0.67 (0.06) |
| | | 0.10 | 0.00 (0.00) | 0.18 (0.00) | 0.70 (0.12) | 0.69 (0.11) |
| | High | 0.05 | 0.00 (0.00) | 0.00 (0.00) | 0.58 (0.05) | 0.59 (0.06) |
| | | 0.10 | 0.00 (0.00) | 0.00 (0.00) | 0.61 (0.11) | 0.61 (0.11) |

Tab. S2: Power comparison between different knockoffs. Multivariate normal features were simulated with $n = 2000$ and $p = 400$, detailed in Section 2. Phenotypes were simulated with 20 driver features whose effect sizes were drawn from standard normal but rescaled to achieve PVE of 0.8. Columns Corr denote feature correlation in simulation: the top half of the table are equi-correlation, and the bottom half are varying correlation with low, medium and high correlations detailed in Tab S1. In each cell we have power and realized FDR (in parenthesis). Columns ReKo-Null and ReKo-Norm are two varieties of Reflection Knockoffs, where ReKo-Null means random component was generated by sampling vectors in the null space while ReKo-Norm means random component was generated by drawing from the standard normal distribution. The prior for reflection is $\sqrt{p}I_p$. The knockoff filter used here is ⊗glmnet⊗Beta.

### 5 Comparison via Multiple Knockoff Filters

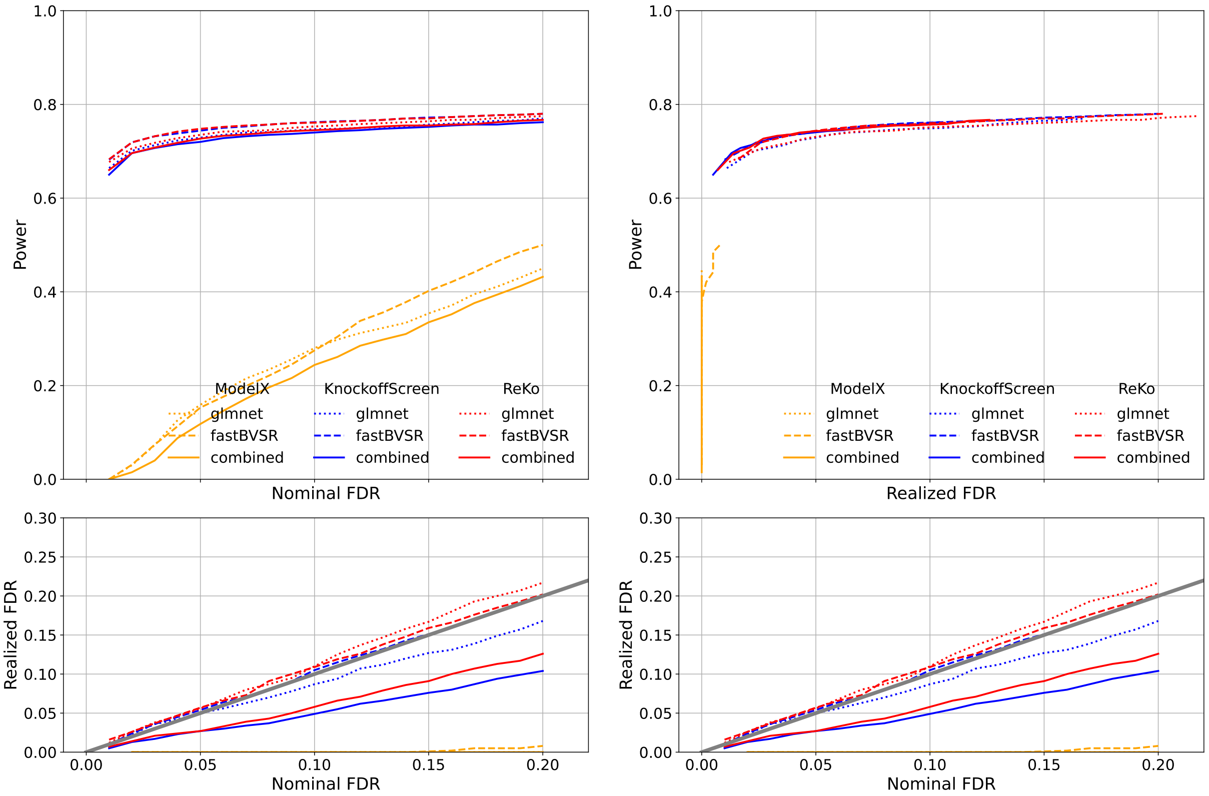


Fig. S2: Power and realized FDR comparison between different knockoffs. Features were simulated following multivariate normal distribution detailed in Section 2 with $n = 2000$, $p = 800$ and medium correlation. The number of driver features is 20 and their effect sizes were drawn from standard normal but rescaled to achieve PVE of 0.8. The results were pooled from 100 replications. For each replicate, 10 knockoff statistics were aggregated to call true and false positives. In left panel, the top plot is nominal FDR vs power and the bottom plot is nominal FDR vs realized FDR. In right panel, the top plot is realized FDR vs power and the bottom plot is identical to the bottom plot in the left panel. Different line styles represent different knockoff filters and different colors represent different knockoff methods.

### 6 Comparison via Multiple Knockoff Filters

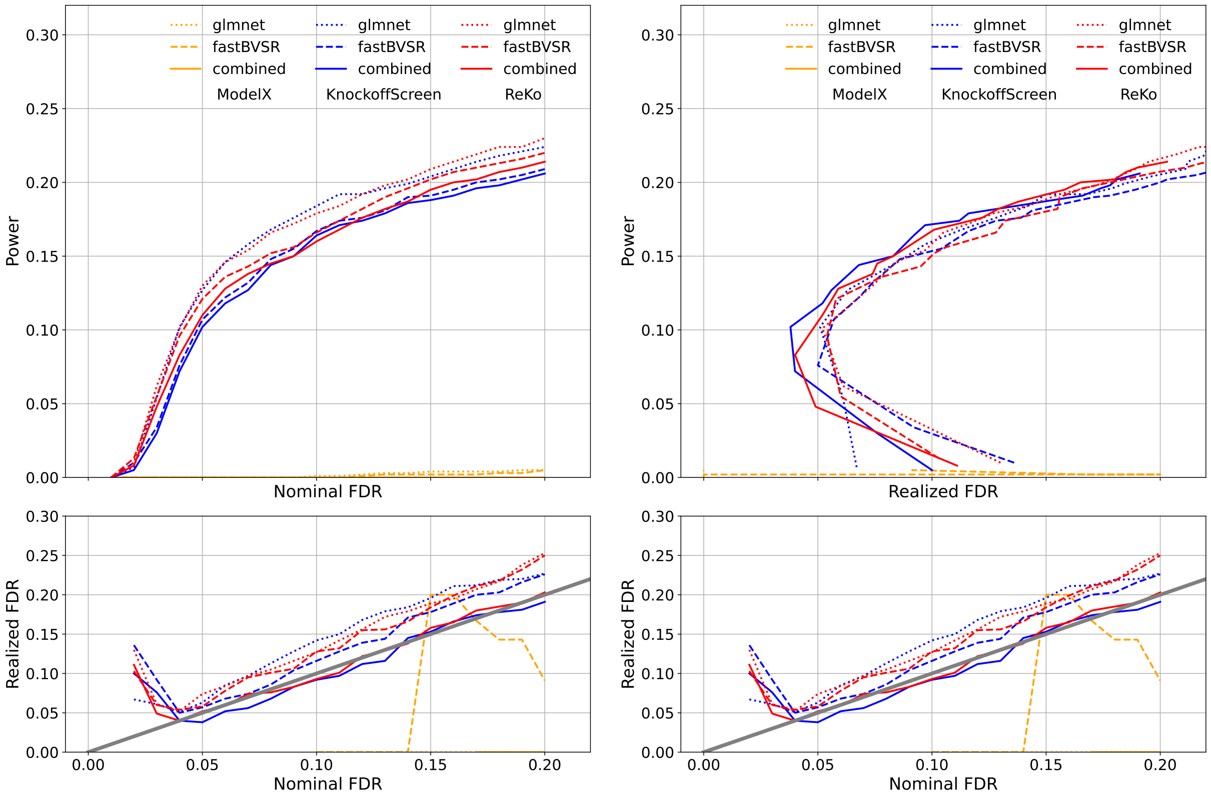


Fig. S3: Power and realized FDR comparison between different knockoffs. Features were simulated following multivariate normal distribution detailed in Section 2 with $n = 2000$, $p = 800$ and medium correlation. The number of driver features is 20 and their effect sizes were drawn from standard normal but rescaled to achieve PVE of 0.2. The results were pooled from 100 replications. For each replicate, 10 knockoff statistics were aggregated to call true and false positives. In left panel, the top plot is nominal FDR vs power and the bottom plot is nominal FDR vs realized FDR. In right panel, the top plot is realized FDR vs power and the bottom plot is identical to the bottom plot in the left panel. Different line styles represent different knockoff filters and different colors represent different knockoff methods. The small PVE appears to produced biased estimates for small nominal FDR for KnockoffScreen and ReKo. Model-X appears to have no power for small PVE.

### 7 Comparison with Multivariate Normal Features

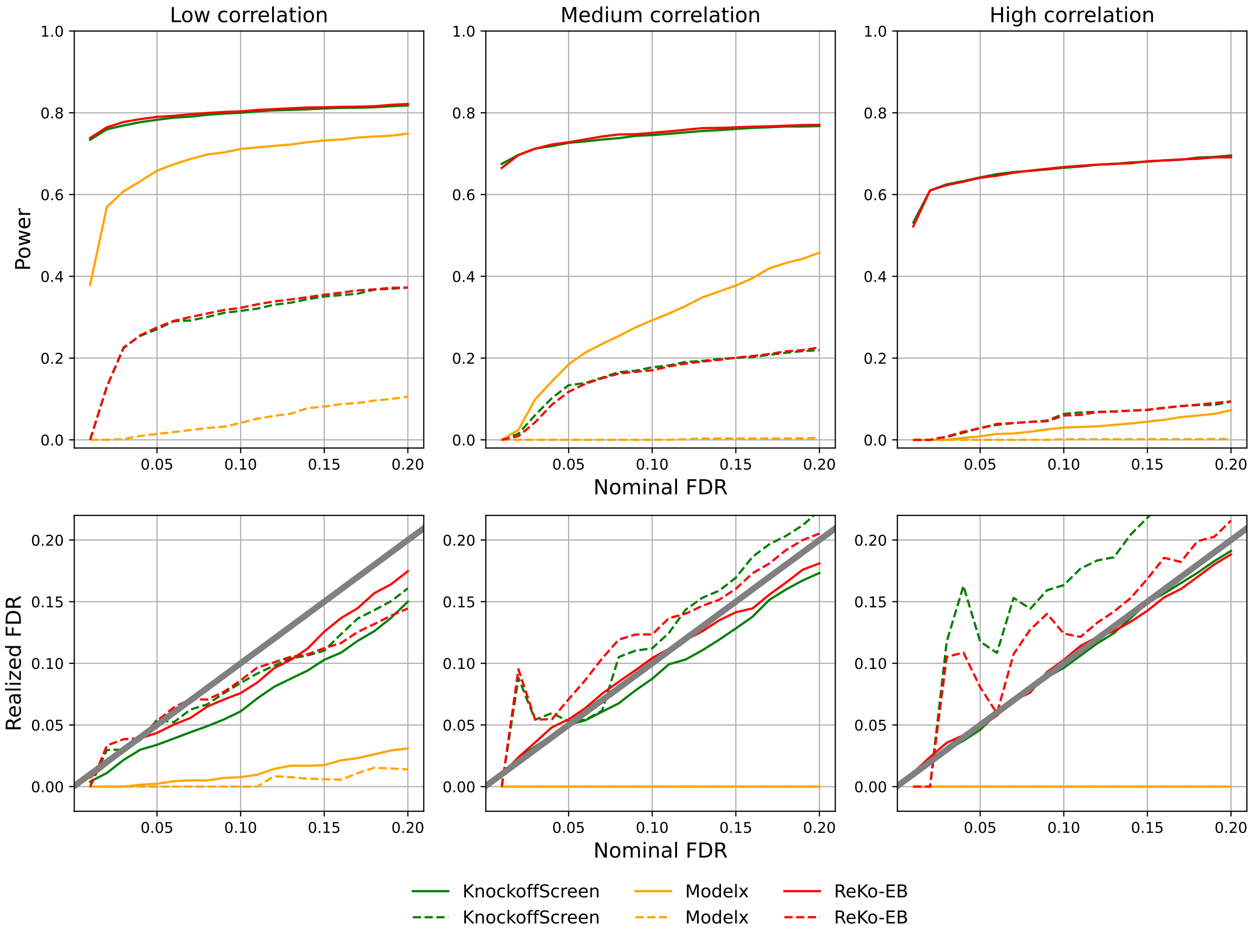


Fig. S4: Power and realized FDR comparison between different knockoffs. Features were simulated following Multivariate Normal Distribution detailed in Section 2. In particular, $n = 2000$, $p = 800$, with low, medium, and high correlations detailed in Fig. S1. Each setting was simulated 100 replications. The number of driver features is 20 and their effect sizes were drawn from standard normal but rescaled to achieve PVE of 0.8 for solid lines and 0.2 for dashed lines. For each simulation setting, 10 knockoff statistics were aggregated to call true and false positives. In these comparisons, the knockoff filter is $\otimes$glmnet. Knockoffs were marked in different colors with orange for Model-X, green for KnockoffScreen, and red for ReKo.

### 8 Comparison with Multivariate T Features

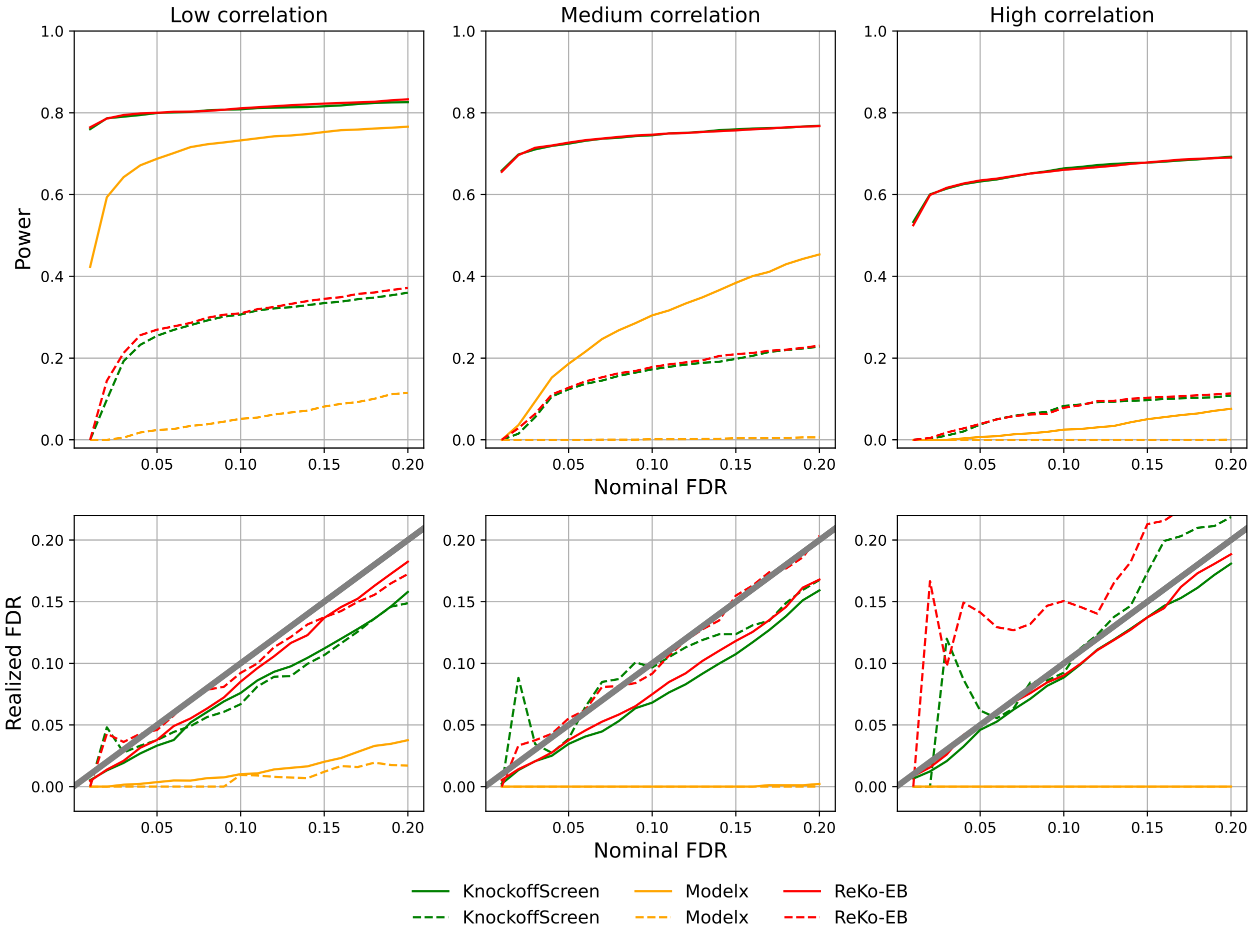


Fig. S5: Power and realized FDR comparison between different knockoffs. Features were simulated following Multivariate T Distribution detailed in Section 2. In particular, $n = 2000$, $p = 800$, with low, medium, and high correlations detailed in Fig. S1. Each setting was simulated 100 replications. The number of driver features is 20 and their effect sizes were drawn from standard normal but rescaled to achieve PVE of 0.8 for solid lines and 0.2 for dashed lines. For each simulation setting, 10 knockoff statistics were aggregated to call true and false positives. In these comparisons, the knockoff filter is ⊗glmnet. Knockoffs were marked in different colors with orange for Model-X, green for KnockoffScreen, and red for ReKo.

### 9 Counts of Positives with and without Aggregation

Counts of positives with aggregation (of 10 sets of knockoff statistics) was documented in Table 1 in main text. The results with nominal FDR 0.10 were reproduced in the first row of the table below. Counts of positives without aggregation were documented in the table below, where the counts were obtained from a single knockoff statistics. The mean and sd are for 10 counts of positives without aggregation.

| n | MX | KS | ReKo |
|---|---|---|---|
| 10 | 308 | 361 | 371 |
| 1 | 350 | 343 | 382 |
| 1 | 277 | 355 | 363 |
| 1 | 315 | 366 | 393 |
| 1 | 290 | 354 | 364 |
| 1 | 321 | 335 | 359 |
| 1 | 310 | 366 | 335 |
| 1 | 290 | 357 | 364 |
| 1 | 258 | 359 | 339 |
| 1 | 302 | 350 | 360 |
| 1 | 321 | 359 | 366 |
| mean | 303.4 | 354.4 | 362.5 |
| sd | 26.0 | 9.7 | 17.2 |

Tab. S3: Counts of significant association for proteomic data at nominal FDR 0.10. Column MX contains results from Model-X⊗fastBVSR; Column KS contains results from KnockoffScreen ⊗combined; Column ReKo contains results from ReKo⊗combined. Column $n$ contains numbers of knockoff statistics aggregated to call positives.

### 10 Speed and Time

We use two example datasets to illustrate the computation time of constructing reflection knockoffs, and model fitting time using different methods. The UK Biobank dataset has $n = 14,752$ (sample) and $p = 1,459$ (feature). Constructing 10 copies of reflection knockoffs takes 14.2 minutes using our C program reko with default options, and takes 3.3 minutes if rSVD approximate computation was invoked (with -b 400). Fitting glmnet takes on average 2.5 minutes; and fitting fastBVSR takes on average 246.6 minutes with $1,000,000$ MCMC steps. Taking *GJB6* as an example we illustrate the computation time for fine mapping. The dataset has $n = 1,477$ and $p = 1,140$. Constructing 10 copies of reflection knockoffs takes 36.4 seconds using our C program reko with default options, and takes only 13.7 seconds if rSVD approximate computation was invoked (with -b 400). Fitting glmnet takes on average 10.5 second; fitting SuSiE takes on average 20.2 seconds; and fitting fastBVSR takes on average 90.0 seconds with $1,000,000$ MCMC steps. Note both reko and glmnet used multithreading (with maximum 112 threads), and SuSiE and fastBVSR used a single thread.
